# Determinants of Spike Infectivity, Processing and Neutralization in SARS-CoV-2 Omicron subvariants BA.1 and BA.2

**DOI:** 10.1101/2022.04.13.488221

**Authors:** Chiara Pastorio, Fabian Zech, Sabrina Noettger, Christoph Jung, Timo Jacob, Konstantin M.J. Sparrer, Frank Kirchhoff

## Abstract

The SARS-CoV-2 Omicron variant rapidly outcompeted other variants and currently dominates the COVID-19 pandemic. Its enhanced transmission, immune evasion and pathogenicity is thought to be driven by numerous mutations in the Omicron Spike protein. Here, we examined the impact of amino acid changes that are characteristic for the BA.1 and/or BA.2 Omicron lineages on Spike function, processing and susceptibility to neutralization. Individual mutations of S371F/L, S375F and T376A in the ACE2 receptor-binding domain as well as Q954H and N969K in the hinge region 1 impaired infectivity, while changes of G339D, D614G, N764K and L981F moderately enhanced it. Most mutations in the N-terminal region and the receptor binding domain reduced sensitivity of the Spike protein to neutralization by sera from individuals vaccinated with the BNT162b2 vaccine or therapeutic antibodies. Our results represent a systematic functional analysis of Omicron Spike adaptations that allowed this SARS-CoV-2 variant to overtake the current pandemic.

**HIGHLIGHTS:**

- S371F/L, S373P and S375F impair Spike function and revert in some BA. 1 isolates
- Changes of Q954H and N969K in HR1 reduce while L981F enhances S-mediated infection
- Omicron-specific mutations in the NTD and RBD of Spike reduce neutralization
- N440K, G446S, E484A and Q493K confer resistance to bamlanivimab or imdevimab

## INTRODUCTION

SARS-CoV-2, the causative agent of the Coronavirus disease 2019 (COVID-19) pandemic, has infected more than 500 million people around the globe and caused almost 6.2 million fatalities (https://coronavirus.jhu.edu/map.html; April 13^th^, 2022). Effective vaccination is the best way to get this devastating pandemic under control. A variety of safe and effective vaccines against SARS-CoV-2 are available and more than 10 billion vaccine doses have been administered to date. However, low access to, or acceptance of vaccines together with the emergence of new SARS-CoV-2 variants jeopardise this strategy. So called variants of concern (VOCs) pose a particular risk. Their increased transmissibility, efficient immune evasion and altered pathogenicity are mainly determined by the viral spike (S) protein (Harvey et al., 2021; Jung et al., 2022; Tao et al., 2021).

Currently, the fifth SARS-CoV-2 VOC, termed Omicron, dominates the COVID-19 pandemic. The Omicron VOC was first detected in Botswana and South Africa in November 2021 and outcompeted the Delta VOC in an amazingly short time. Evolutionary studies revealed that the Omicron VOC evolved independently, possibly in a chronically infected immunocompromised individual, human population under poor surveillance or an unknown non-human species from which it spilled back to humans (Karim et al., 2021; Wei et al., 2021). Omicron contains a strikingly high number of mutations (Jung et al., 2022), especially in its S protein, compared to other variants and the initial Wuhan strains. Recent studies support that this VOC is highly transmissible and shows an increased ability to infect convalescent and vaccinated individuals (Altarawneh et al., 2022; Espenhain et al., 2021; Grabowski et al., 2022; Pulliam et al., 2022). This agrees with the finding that the Omicron VOC shows reduced susceptibility to neutralizing antibodies induced by previous SARS-CoV-2 infection or vaccination (Andrews et al., 2021; Cele et al., 2021; Hoffmann et al., 2021; Lu et al., 2021; Planas et al., 2021; Wilhelm et al., 2021). Notably, accumulating evidence suggests that Omicron infections are associated with milder symptoms and decreased hospitalization and fatality rates compared to infections with the Delta SARS-CoV-2 VOC (Moore and Baden, 2022; Wolter et al., 2022).

The SARS-CoV-2 S protein is the major membrane glycoprotein required for recognising the viral receptor angiotensin-converting enzyme 2 (ACE2) and subsequent entry into target cells (Hoffmann et al., 2020; Letko et al., 2020). Thus, the S protein critically determines the cell tropism and transmissibility of SARS-CoV-2 in human populations. To mediate attachment and fusion, the S precursor needs to be proteolytically processed by cellular proteases after synthesis. The proprotein convertase furin cleaves S at the S1/S2 site to generate the S1 subunit, which is responsible for receptor binding, while the transmembrane serine protease 2 (TMPRSS2) or cathepsins B and L cleave at the S2’ site just upstream of the hydrophobic fusion peptide to release the S2 subunit mediating membrane fusion (Walls et al., 2020; Wrapp et al., 2020). In its active form the S protein of SARS-CoV-2 forms trimers on the surface of the viral particles. Consequently, the S protein is also the major target of protective humoral immune responses (Walls et al., 2020) and all COVID-19 vaccines are based on the SARS-CoV-2 S antigen. Thus, mutations in the N-terminal or receptor binding domains (NTD and RBD, respectively) of S can confer increased resistance to neutralizing antibodies (Dai and Gao, 2021). It is clear that alterations in the viral surface glycoprotein Spike (S) of the Omicron VOC play a key role in its high transmissibility, efficient immune evasion and reduced pathogenicity. However, the impact of most amino acid changes distinguishing the Omicron S protein from that of the original Wuhan SARS-CoV-2 strain on viral infectivity and susceptibility to neutralization remains to be determined.

Here, we analyzed the functional impact of individual amino acid changes that distinguish the dominant 21K (BA.1) Omicron VOC and the emerging 21L (BA.2) variant from the early 2020 Wuhan SARS-CoV-2 isolate. To achieve this, we introduced a total of 48 mutations in the S protein of the Wuhan strain and determined their impact on viral infectivity, expression and proteolytic processing, as well as susceptibility to neutralizing antibodies and sera from vaccinated individuals. We show that several amino acid changes found in the Omicron S protein impair infectivity and demonstrate that numerous alterations in the NTD and RBD of BA.1 and or BA.2 S proteins affect neutralization by sera from BNT/BNT vaccinated individuals and therapeutic antibodies.

## RESULTS

### Generation of S Proteins containing mutations found in Omicron

Omicron is currently classified into two major lineages, BA.1 (21K) and BA.2 (21L) (Figure 1A) (Hadfield et al., 2018; Sagulenko et al., 2018). BA.1 has replaced the Delta VOC and dominated the COVID-19 pandemic at the beginning of 2022 (Figure 1B). However, the frequency of the BA.2 lineage is increasing (Figure 1B) and this variant has outcompeted BA.1 in many countries, such as India, Denmark, Austria and South Africa (Viana et al., 2022). Although only ~13% of the SARS-CoV-2 genome encodes for the S protein, this region contains most mutations distinguishing the Omicron VOCs from the original Wuhan Hu-1 SARS-CoV-2 strain. Many of the mutations that distinguish the Omicron S proteins from those of other SARS-CoV-2 variants are located in the RBD that interacts with the viral ACE2 receptor and is a major target of neutralizing antibodies (Figure 1C). The Omicron BA.1 and BA.2 S proteins share about 20 amino acid changes in S compared to the 2020 Wuhan Hu-1 strain and 12 of these are located in the RBD (Figures 1C, 1D). In the consensus, a total of 14 mutations are specific for BA.1 and 9 for BA.2 (Figures 1C, 1D). Thirteen of the 23 S consensus lineage-specific variations are located in the NTD (Figure 1D). All 43 non-synonymous defining mutations, insertions and deletions found in BA.1 and BA.2 Omicron VOCs (https://covariants.org/variants/21L) were introduced individually in the S protein of the original Wuhan Hu-1 strain by site-directed mutagenesis. Sequence analysis of the full-length S genes verified that all constructs contained the desired mutations (Figure 1D) and confirmed the absence of additional changes.

**Figure 1.**
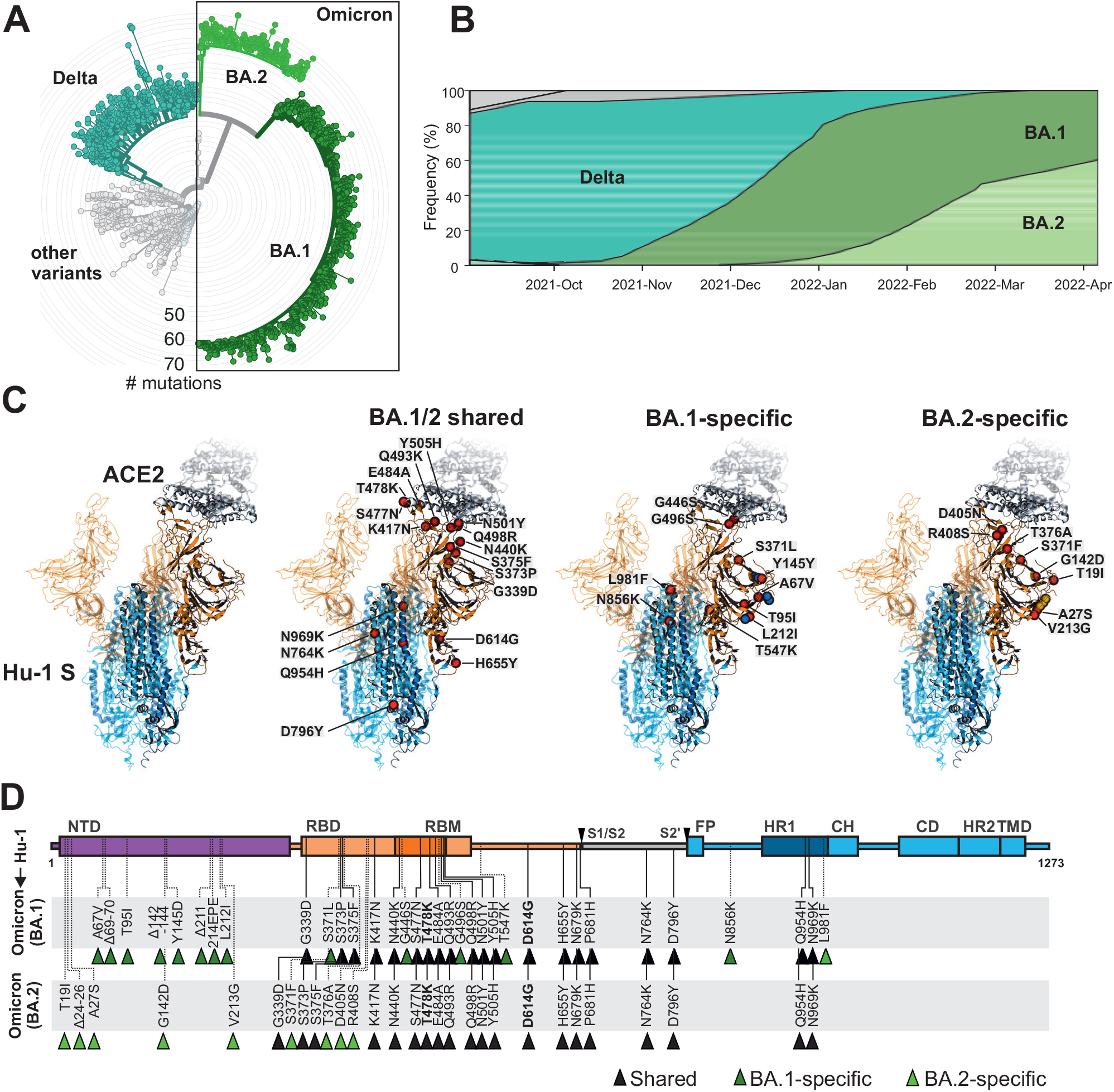
Features of Omicron BA.1 and BA.2 VOCs. (A)Radial phylogenetic tree of representative SARS-CoV-2 strains (n= 2863 genomes, sampled between Dec 2019 and Mar 2022) scaled according to their divergence compared to the Wuhan Hu-1 sequence. Retrieved from Nextstrain on April 7^th^ 2022 (https://nextstrain.org/ncov/gisaid/global?m=div) and modified. Colour coding according to VOCs as indicated. (B) Frequencies of SARS-CoV-2 Delta, BA.1 and BA.2 sequences in data from GenBank from September 1^st^ 2021 to 7^th^ of April 2022. Scaled to 100%. Retrieved and modified from Nextstrain on April 7^th^ 2022. Orange, Delta VOC. Purple, BA.1. Green, BA.1. (C) Overview on the SARS-CoV-2 spike structure (downloaded from PDB: 7KNB) and localization of amino acid changes that are shared between BA.1 and BA.2 or specific for BA.1 or BA.2 as indicated. S1 (orange), S2 (blue) ACE2 (grey), mutations (red), BA.1 specific deletions (blue), BA.2 specific deletions (yellow). (D) Schematic depiction of the SARS-CoV-2 spike, its domains and amino acid alterations in Omicron BA.1 (dark green) and BA.2 (light green) VOC compared to the Wuhan Hu-1 sequence. S1 subunit: N-terminal domain, NTD (purple) and receptor binding domain, RBD (orange). Receptor binding motif, RBM (dark orange). S2 subunit: fusion peptide, FP (blue), heptad repeat 1, HR1 (dark blue) central helix, CH, connector domain, CD, heptad repeat 2, HR2, and transmembrane domain, TM (blue).

### Impact of mutations in Omicron Spike on viral pseudoparticle infection

To analyse the functional impact of mutations found in the Omicron BA.1 and BA.2 variants, we generated vesicular stomatitis virus (VSV) particles pseudotyped with the parental and mutant SARS-CoV-2 S proteins. Previous studies established that these VSV pseudoparticles (VSVpp) mimic key features of SARS-CoV-2 entry, such as receptor usage, cell tropism, protease dependency and susceptibility to neutralizing antibodies (Hoffmann et al., 2020; Riepler et al., 2020; Schmidt et al., 2016). We found that the BA.1 S showed moderately reduced and the BA.2 S slightly enhanced infection efficiencies compared to the Wuhan Hu-1 S, while the S protein of the Delta VOC did not differ significantly from the original form (Figure 2A, left). Most of the 20 amino acid changes in S that are shared between the BA. 1 and BA.2 variants did not significantly affect the efficiency of VSVpp infection (Figures 2A; S1). In agreement with previous findings (Korber et al., 2020; Yurkovetskiy et al., 2020), substitution of D614G slightly enhanced infection. Similarly, mutations of G339D, K417N and N764K had subtle enhancing effects. Notably, modest enhancing effects were not due to saturation of infection since only up to 10% of all target cells became GFP positive during the single round of infection (Figure S1). Substitution of S375F in the RBD drastically impaired and mutations of Q954H and N969K in HR1 reduced VSVpp infectivity (Figure 2A; S1).

**Figure 2.**
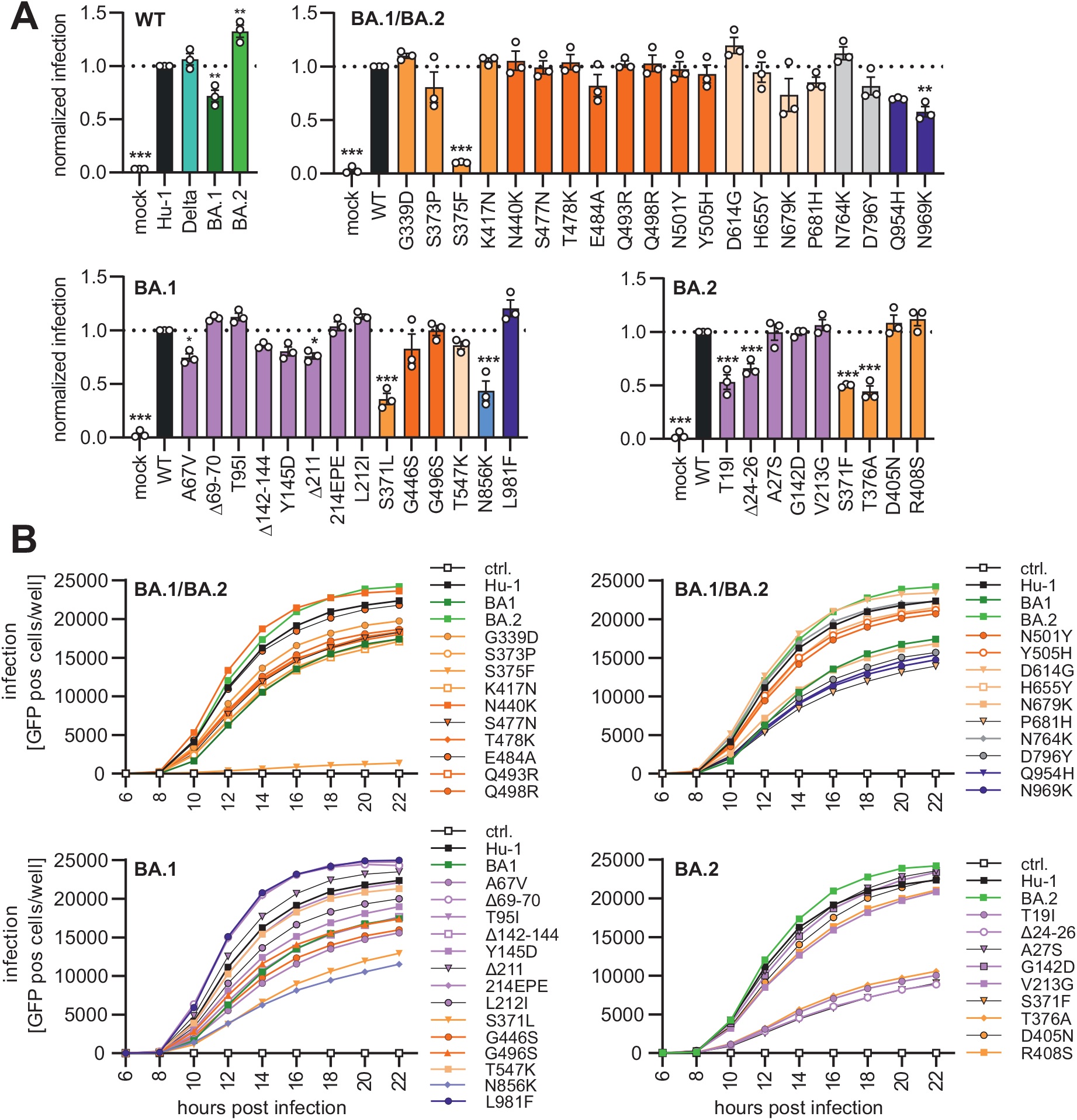
Impact of mutations in Omicron on Spike-mediated infection. (A) Automatic quantification of infection events of CaCo-2 cells transduced with VSVΔG-GFP pseudotyped with SARS-CoV-2 Hu-1 (grey), Delta (turquise), BA.1 (dark green), BA.2 (light green) or indicated mutant S proteins. The localisation of each mutation in S is indicated by colour. S1: NTD (purple), RBD (orange), RBM (dark orange), others (light orange), S2: HR1 (dark blue) others (light blue). Bars represent the mean of three independent experiments (±SEM). Statistical significance was tested by one-way ANOVA. *P <0.05; **P <0.01; ***P <0.001. (B) Infection kinetics of CaCo-2 cells infected by VSVpp containing the indicated mutant S proteins. Infected cells were automatically quantified over a period of 22 h. See also Figure S1.

Most of the BA.1 and BA.2 specific variations in the NTD of the S protein had minor effects on VSVpp infectivity (Figure 2A). Three changes (Δ69-70, T95I and L212I) in the NTD slightly enhanced and six alterations (T19I, Δ24-26, A67V, Δ142-144, Y145D, and Δ211) reduced VSVpp infection. Similar to the shared S375F, mutations of S371L or S371F in the BA.1 and BA.2 S proteins, respectively, strongly impaired viral infectivity. The adjacent BA.2-specific T376A change had similar disruptive effects (Figure 2A). Mutation of N856K that is specific for BA.1 and might stabilize the fusion peptide proximal region (Zhang et al., 2022) and T19I or Δ24-26 near the N-terminus of BA.2 S markedly reduced VSVpp infection (Figure 2A), although these residues do not affect known functional domains.

To assess infection kinetics and to challenge the above-mentioned infection results, we performed assays allowing automated quantification of the number of VSVpp infected (GFP+) Caco-2 cells over time. The various mutant S proteins mediated infection with similar kinetics but varying and frequently reduced efficiencies (Figure 2B). The results confirmed that the BA.1 S shows moderately diminished infection efficiency compared to the Hu-1 Wuhan S protein. Individual mutations of T19I, Δ24-26, A67V, Y145D, S371L, S371F, S373P, S375F, T376A, G446S, Q493R, G496S, N679K, P681H, D796Y, N856K, Q954H and N969K all reduced the activity of the Hu-1 S to levels similar or below that obtained for the BA.1 S protein (Figure 2B). In contrast, shared mutations of N440K and D614G, as well as BA.1-specific changes of Δ69-70, Δ211, insertion of 214EPE, and mutation of L981F increased infection efficiencies. Our results agree with recent findings suggesting that the Q954H and N969K changes in heptad repeat 1 (HR1) reduce rather than enhance fusion efficiency (Suzuki et al., 2022; Xia et al., 2022; Zhao et al., 2021). In addition, our analysis revealed that N856K in BA.1 S and T19I as well as Δ24-26 in the BA.2 NTD strongly impaired S-mediated infection. Perhaps most notably, all individual mutations in the three serine residues in a small loop region (S371L/F, S373P, S375F), as well as the adjacent BA.2-specific T376A change severely impaired S-mediated infection.

### Inefficient Processing and Virion Incorporation of Specific Spike Variants

To examine S expression, proteolytic processing and virions incorporation of the mutants, we performed comprehensive western blot analyses of HEK293T cells co-transfected with VSVΔG-eGFP and S expression constructs and the S-containing VSVpp in the culture supernatants. In agreement with the infectivity data, most individual amino acid changes, deletions or insertions had no significant impact on S expression and processing (Figure 3A). All 44 parental and mutant full-length S proteins were readily detected in the cellular extracts (Figure 3A). However, mutations in S371L, S373P, S375F and T376A that impaired S infectivity (Figure 2) also reduced the efficiency of processing and/or incorporation into viral pseudoparticles (Figure 3A). The phenotypes of the S375F and T376A mutant were most striking and these S variants were hardly processed. Two BA.2 specific alterations in S (T19I and Δ24-26) that were less active in infection assays were associated with reduced levels of S protein on VSVpp (Figure 3A). Altogether, the levels of S2 protein expression and processing in cellular extracts relative to the parental Hu-1 S proteins correlated well with one another (Figure S2A) and with the efficiency of S-mediated infection (Figure 3B, left). Similar but less significant correlations were observed for the VSVpp infection and Spike levels in the culture supernatants (Figure 3B, right). T19I, Δ24-26, S375F and T376A reduced the levels of both S and S2 incorporated into VSVpp, while S371L/F mainly affected S2 levels in the particles. In comparison, mutations of Q493R, T547K, D796Y and N856K reduced VSVpp infection without exerting significant effects on S expression and processing in the cells although T547K and D796Y were associated with reduced levels of S2 in VSVpp (Figure 3). None of the mutations (H655Y, N679K and P681H) located near the S1/S2 cleavage site had significant effects on S processing (Figure 3). In addition, confocal microscopy showed that mutant S proteins showing enhanced (D614G, L981F) or impaired (T19I, S371L/F, S373P, S375F, T376A) activity all localized at the cell surface (Figure S2B) indicating that disruptive effects were not due to impaired trafficking or mislocalization. Altogether, our results revealed that changes of T19I, Δ24-26, T376A, S375F and Q954H reduce VSVpp infectivity by affecting S processing although they are not located in proximity to the S1/S2 furin cleavage site.

**Figure 3.**
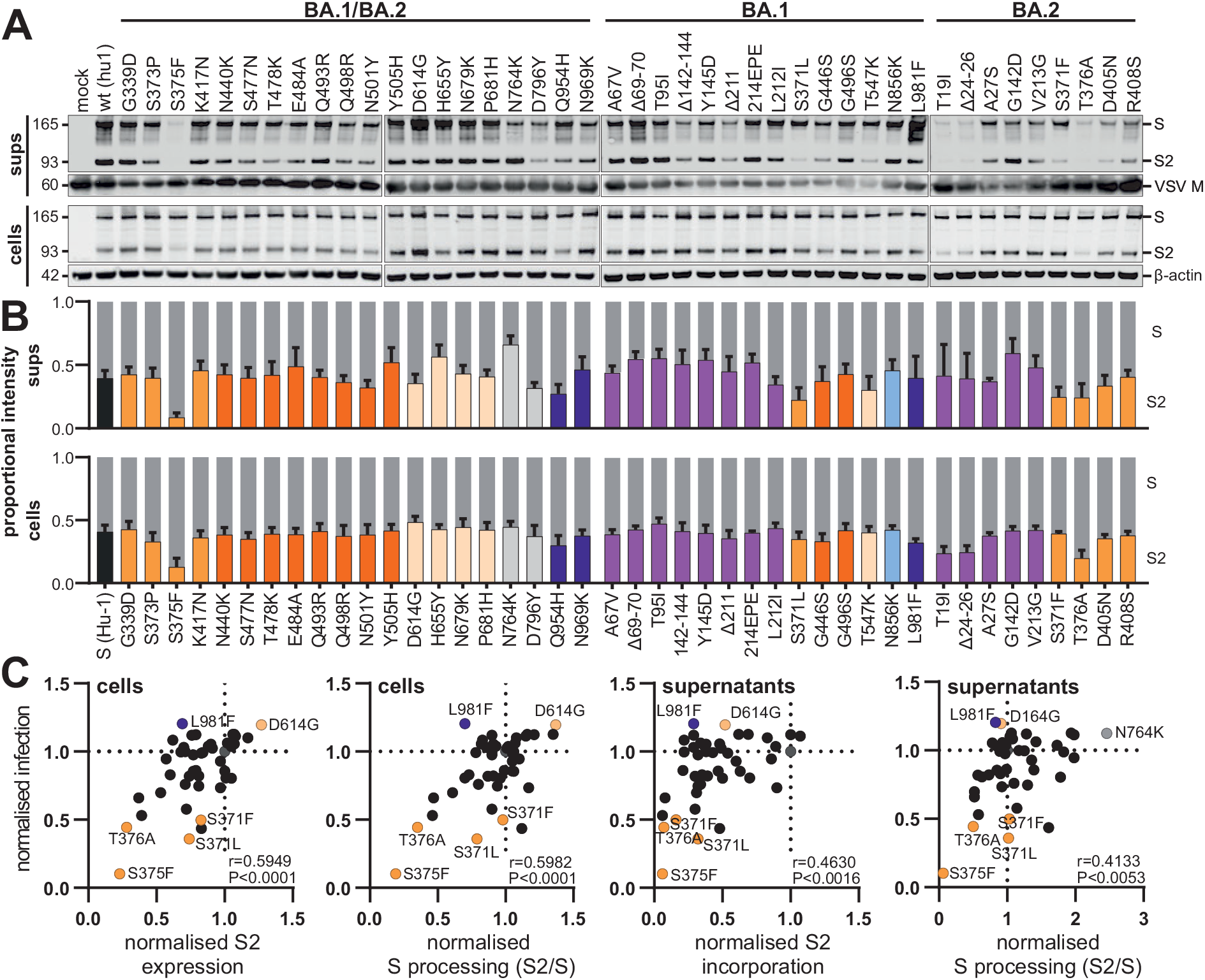
Expression and processing of Spike proteins containing mutations present in the Omicron BA.1 and BA.2 VOCs. (A) The upper panels show exemplary immunoblots of whole cells lysates (WCLs) and VSVpp containing supernatants of HEK293T cells transfected with vectors expressing the Hu-1 or mutant SARS-CoV-2 S proteins and VSVΔG-GFP constructs. Blots were stained with anti-V5 (Spike), anti-ß-actin and anti-VSV-M protein. Lower panels: expression levels of uncleaved, full-length Spike protein (S, gray bars) and the S2 subunit (bars coloured according to the corresponding domains as shown in Figure 1D) were quantified. The results show mean values (±SEM) obtained from three independent experiments. (B) Correlation of the S2 expression/incorporation and S/S2 processing of the parental S Hu-1 or indicated mutant S proteins in cells and supernatants with the corresponding pseudotype infection rates. Coefficient of determination (R^2^-values) and two tailed P values are provided. See also Figure S2.

### Functional relevance of serine mutations in an RBD loop region

It came as surprise that all individual mutations of S371, S373 and S375 that are found in the Wuhan Hu-1 strains and the Alpha, Beta, Gamma and Delta VOCs to 371L/F, 373P and 375F present in Omicron severely impaired S function. Analysis of available SARS-CoV-2 sequences revealed that the BA.1 and BA.2 S proteins usually contain combined changes of SxSxS to FxPxF or LxPxF, respectively (Figure 4A). However, we identified a small subcluster within the BA.1 sequence showing reversions to serine residues. Phylogenetic analysis suggests that these occurred sequentially: from FxPxF to FxSxF to FxSxS to SxSxS (Figure 4A). The serine containing loop is located adjacent to the RBD and might affect its up and down state (Sztain et al., 2021) and stabilize RBD-RBD interactions (Wrobel et al., 2022) (Figure 4B). Altogether, the results suggested that the combination of S371-S373-S375 or 371F/L373P375F might be required for effective S function and processing. To address this experimentally, we generated the LPF (BA.1) and FPF (BA.2) triple mutants of the Hu-1 S protein and analysed their ability to mediate VSVpp infection. Both showed dramatically lower fusion activity (Figure 4C). In comparison, combined changes of S477N/T478K in the RBD and N764K/N856K/Q954H in S2 had only modest disruptive effects and alterations of N679K/P681H near the S2’ processing site did not significantly change the infection efficiency of the Hu-1 S protein (Figure 4C). Intracellular localisation analyses showed that the LxPxF and FxPxF mutant S proteins were readily detectable at the cell surface, just like the parental Hu-1 S protein (Figure 4D). Thus, in agreement with the results on individual mutations (Figure S2B) the impaired activity and processing of the triple mutant S proteins is not due to altered trafficking or subcellular localization.

**Figure 4.**
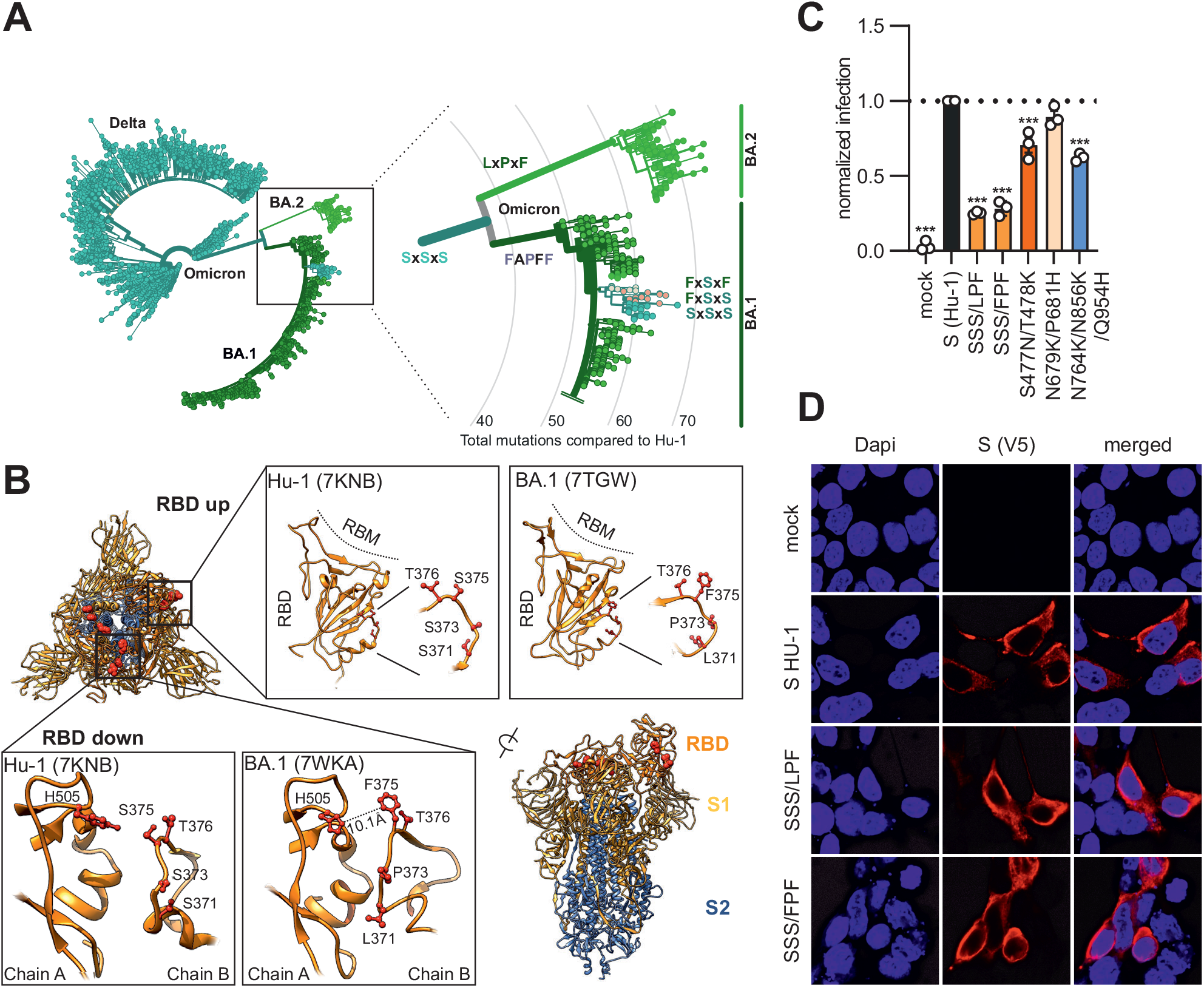
Evolution and functional relevance of S371L/F, S373P and S375F changes in the Spike protein. (A) Phylogenetic tree of Delta, Omicron BA.1 and BA.2 strains. Amino acids at position 371, 373 and 375 are indicated; all other SARS-CoV-2 variants almost invariantly contain three serines at these positions. Colour coding as indicated according to VOC. Retrieved and modified from Nextstrain on April 7^th^ 2022. (B) Close-up view of the region encompassing the mutations S371L, S373P and S375F and the surrounding region. Downloaded from PDB: 7KNB, 7TGW or 7WKA as indicated. (C) Automatic quantification of infection events of CaCo-2 cells transduced with VSVΔG-GFP pseudotyped with SARS-CoV-2 Hu-1 or indicated combined mutations. Bars represent the mean of three independent experiments (±SEM). Statistical significance was tested by one-way ANOVA. *P <0.05; **P <0.01; ***P <0.001. (D) Immunofluorescence images of HEK293T cells expressing the parental S Hu-1, the BA.1 specific S37xLPF or the BA.2 specific S37xFPF mutations. Scale bar, 10 μm.

### Effect of Mutations in Omicron Spike on ACE2 interaction and Cell-to-Cell Fusion

To examine the impact of specific mutations in the Omicron S protein on ACE2 interaction, we used a previously established *in vitro* S-ACE2 binding assay (Zech et al., 2021). Immobilized ACE2 is incubated with lysates of transfected HEK293T cells transfected with mutant S expression constructs. S protein retained after washing is detected by an αV5-Ms and quantified using a secondary HRP-conjugated anti-mouse Ab. The S371F, S373P, D614G, N856K and L981F mutations in the Hu-1 S had little if any effect on S binding to human ACE2 (Figure 5A). In comparison, individual substitutions of S375F and T376A and the triple mutations (SSS to LPF or FPF) reduced the levels of S protein bound to ACE2 (Figure 5A). In line with published data (Tian et al., 2011), mutation of N501Y enhanced binding of the SARS-CoV-2 S protein to human ACE2 (Figure 5A).

**Figure 5.**
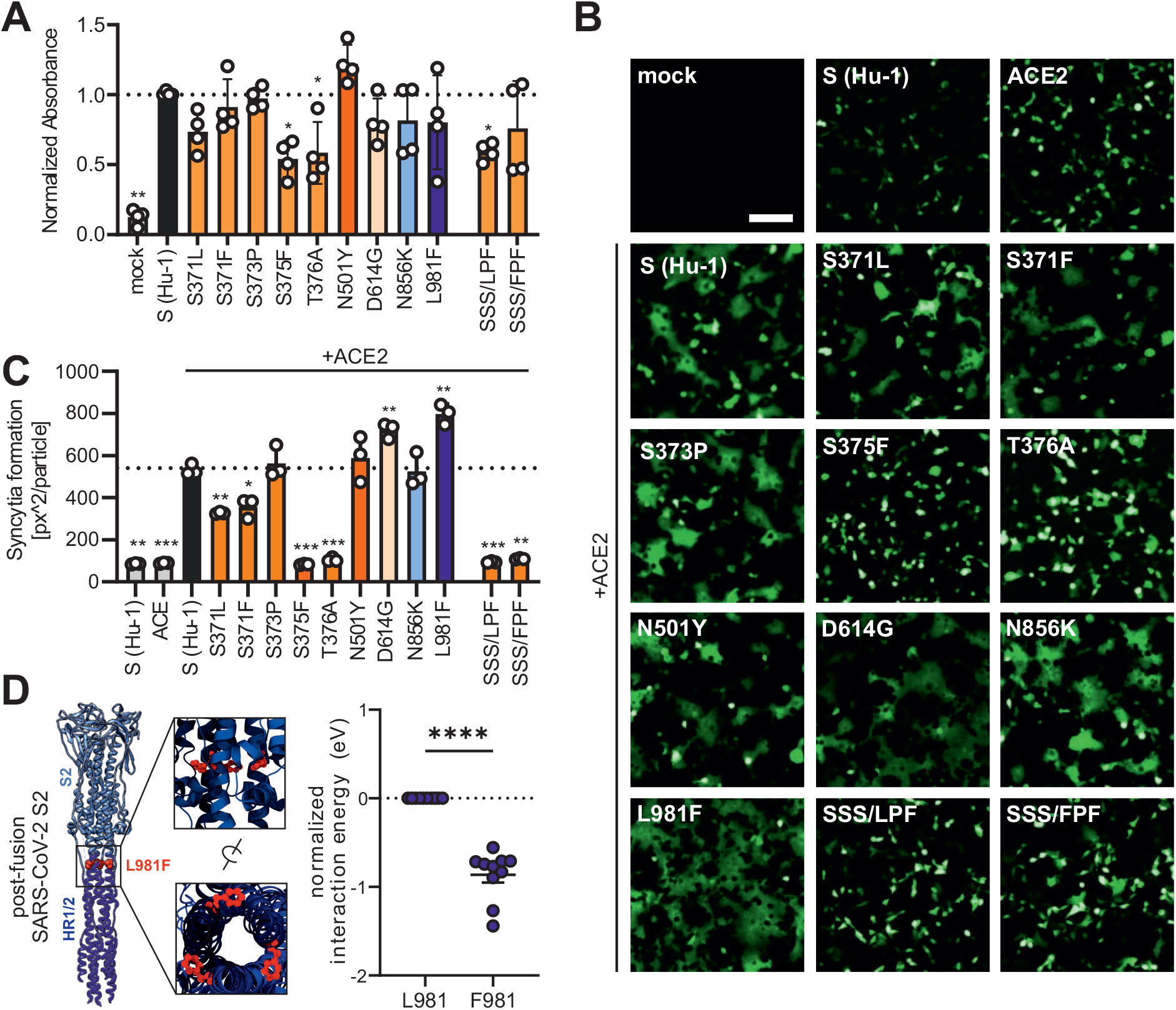
ACE2 interaction of Spike variants. (A) Binding of the indicated Hu-1 and mutant S proteins to ACE2 binding using whole cell lysates of transfected HEK293T. Bars represent the mean of three independent experiments (±SEM). Statistical significance was tested by one-way ANOVA. *P <0.05; **P <0.001. (B) Representative fluorescence microscopy images of HEK293T cells expressing parental Hu-1 or indicated mutant S proteins, Human ACE2 and GFP (green). Scale bar, 125 μm. (C) Automatic quantification of syncytia formation of HEK293T cells expressing parental Hu-1 or indicated mutant S proteins and Human ACE2. Bars represent the mean of three independent experiments (±SEM). Statistical significance was tested by two-tailed Student’s t-test with Welch’s correction. *P <0.05; **P <0.01; ***P <0.001. (D) Overview on the SARS-CoV-2 post-fusion spike structure (downloaded from PDB: 6M3W) and comparative ReaxFF simulation of the mutation L981F.

Cell-to-cell fusion assays showed that co-expression of human ACE2 and the parental Hu-1 as well as the S373P, N501Y, D614G, N856K and L981F S proteins resulted in the formation of large syncytia (Figure 5B, 5C). In contrast, the S375F, T376A and triple LxPxF or FxPxF mutant S proteins did not lead to detectable fusion, while intermediate phenotypes were observed for the S371L and S371F Spikes (Figures 5B, 5C). In agreement with the VSVpp infection data, these results show that individual or combined mutations in S371, S373 and S375 or T376 disrupt the ability of the S protein to mediate membrane fusion.

Mutations of D614G and (to a stronger extent) L981F significantly increased syncytia formation (Figures 5B, 5C). L981 is located in the HR1 region of the S2 protein that interacts with HR2 to form a six-helix bundle to drive virus–host or cell-to-cell membrane fusion (Figure 5D). In agreement with the functional data, molecular modelling of HR1/HR2 interactions using reactive force field simulations predicted that mutation of L981F significantly enhances interactions between HR1 and HR2 (Figure 5D). Taken together, syncytia formation is promoted by D614G found in all VOCs and the Omicron-specific mutation L981F, but almost abrogated by S375F, T376A and the triple SSS to LPF or FPF changes.

### Mutations in the Omicron S Affect Neutralization by Sera from Immunized Individuals

Recent studies have shown that the Omicron BA.1 and BA.2 Spikes show reduced sensitivity to neutralizing Abs induced upon infection and vaccination (Andrews et al., 2021; Cele et al., 2021; Hoffmann et al., 2021; Lu et al., 2021; Zhang et al., 2021). To determine the contribution of individual amino acid changes to immune evasion by Omicron, we compared the sensitivity of the four parental Hu-1, Delta, BA.1, BA.2 with 43 mutant S proteins, each harboring one Omicron-specific mutation, to neutralization by sera from five individuals who received a prime boost vaccination with the mRNA-based BioNTech-Pfizer (BNT162b2) vaccine (Table S1). This vaccine has been approved in 141 countries (https://covid19.trackvaccines.org/vaccines/6/), is frequently used in Europe and the US, and induces efficient protection against most COVID-19 variants (Polack et al., 2020) but shows about 5- to 40-fold lower efficiency against Omicron (Cele et al., 2021; Collie et al., 2022; Iketani et al., 2022; Lu et al., 2021). Predictably, five randomly selected sera collected two weeks after the second dose of BNT neutralized BA.1 and BA.2 on average with substantially lower efficiency than the original Wuhan Hu-1 and Delta variants (Figure 6A; Table S1). A variety of shared as well as BA.1 or BA.2 specific amino acid changes reduced sensitivity to neutralization (examples shown in Figure 6A). It has been previously shown that the NTD contains important neutralizing epitopes (Chi et al., 2020) and that the mutations, deletions and insertions in this region are associated with significant structural changes (Zhang et al., 2022). Our analyses revealed that most individual mutations found in the NTD of BA.1 and BA.2 S proteins reduced neutralization sensitivity (Figure 6B). Deletion of residues 142-144 in BA.1 and G142D in BA.2 had the strongest effects (~9-fold reduction) followed by mutation of Y145D and 214EPE (both in BA.1) that conferred ~7-fold resistance. Amino acid changes in the RBD, such as G339D, S371L, S373P, K417N and N440K, as well as BA.2-specific alterations of S371F and R408S reduced sensitivity to neutralization by BNT/BNT sera, usually in the range of ~2- to 5-fold (Figure 6B). In comparison, five of the six mutations in the S2 region had little if any effect on neutralization. Only the N764K change reduced it on average about 2-fold. Altogether, 27 of the 43 mutations analyzed enhanced antibody-mediated neutralization resistance by >2-fold (Figure 6B). This further supports that a large number of substitutions in the Omicron Spike cooperates to allow efficient viral evasion of humoral immune responses.

**Figure 6.**
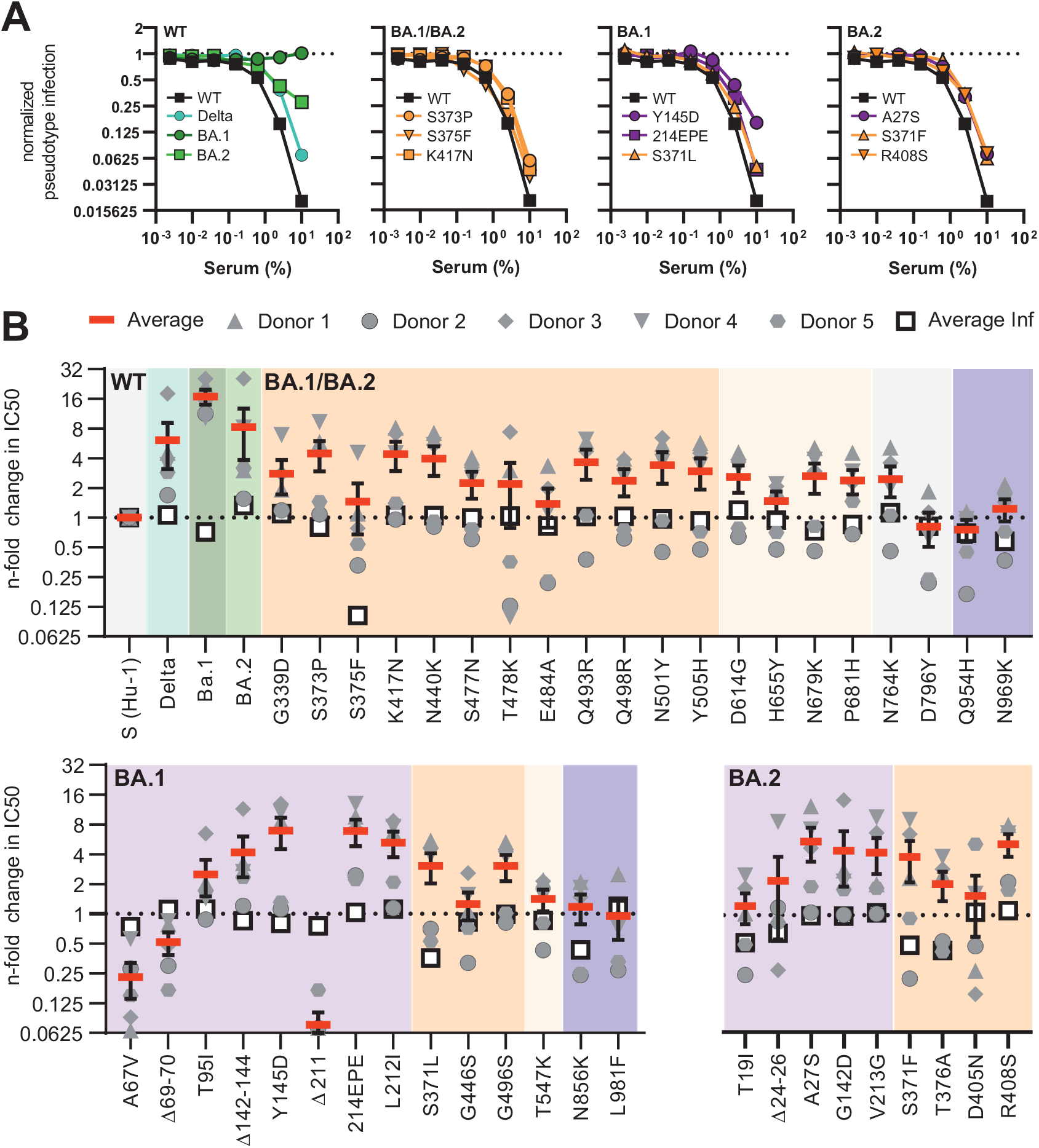
Impact of mutations in the Omicron Spike on serum neutralization. (A) Neutralization of VSVpp carrying the indicated wildtype and mutant S proteins by sera obtained from five BNT/BNT vaccinated individuals compared to the untreated control (set to one). Shown are mean values obtained for the five sera, each tested in three independent experiments. (B) Changes in TCID_50_ values obtained for neutralization of the indicated mutant S proteins by sera from five vaccinated individuals relative to those obtained for the Hu-1 S. Solid red bars indicate average values (±SEM) for the five sera and open black squares the average infectivity of the respective S containing VSVpp shown in Figure 2A. See also Figure Table S1.

In the final set of experiment, we examined the impact of specific mutations in the Omicron S protein on neutralization sensitivity to the FDA-approved therapeutic monoclonal antibodies REGN10987 (marketed as imdevimab), LY-CoV555 (marketed as bamlanivimab) and REGN10933 (marketed as casivirimab). The BA.1 VOC was not inhibited by imdevimab and the N440K or G446S mutations in the Hu-1 S were sufficient to confer full resistance (Figure 7). In contrast, BA.2 S remained susceptible to imdevimab and changes of E484A, Q493R and G496S had little effect. In comparison, both BA.1 and BA.2 were fully resistant to bamlanivimab and substitutions of E484A or Q493R were sufficient to confer resistance (Figure 7). These results agree with those of two recent studies that also examined the impact of individual amino acid changes found in the BA.1 and BA.2 spikes on neutralization by a panel of monoclonal antibodies (Iketani et al., 2022; Liu et al., 2022). Finally, casivirimab showed no appreciable activity against BA.1 but neutralized BA.2 and all mutant S proteins analyzed, albeit with lower efficacy compared to the original Hu-1 S (Figure 7). Altogether, our results show that a strikingly high number of amino acid changes in the NTD and RBD regions of the Omicron S proteins contribute to evasion from neutralizing antibodies. The impact of individual mutations on susceptibility to neutralization varies strongly between sera obtained from individuals who received the BNT/BNT vaccine.

**Figure 7.**
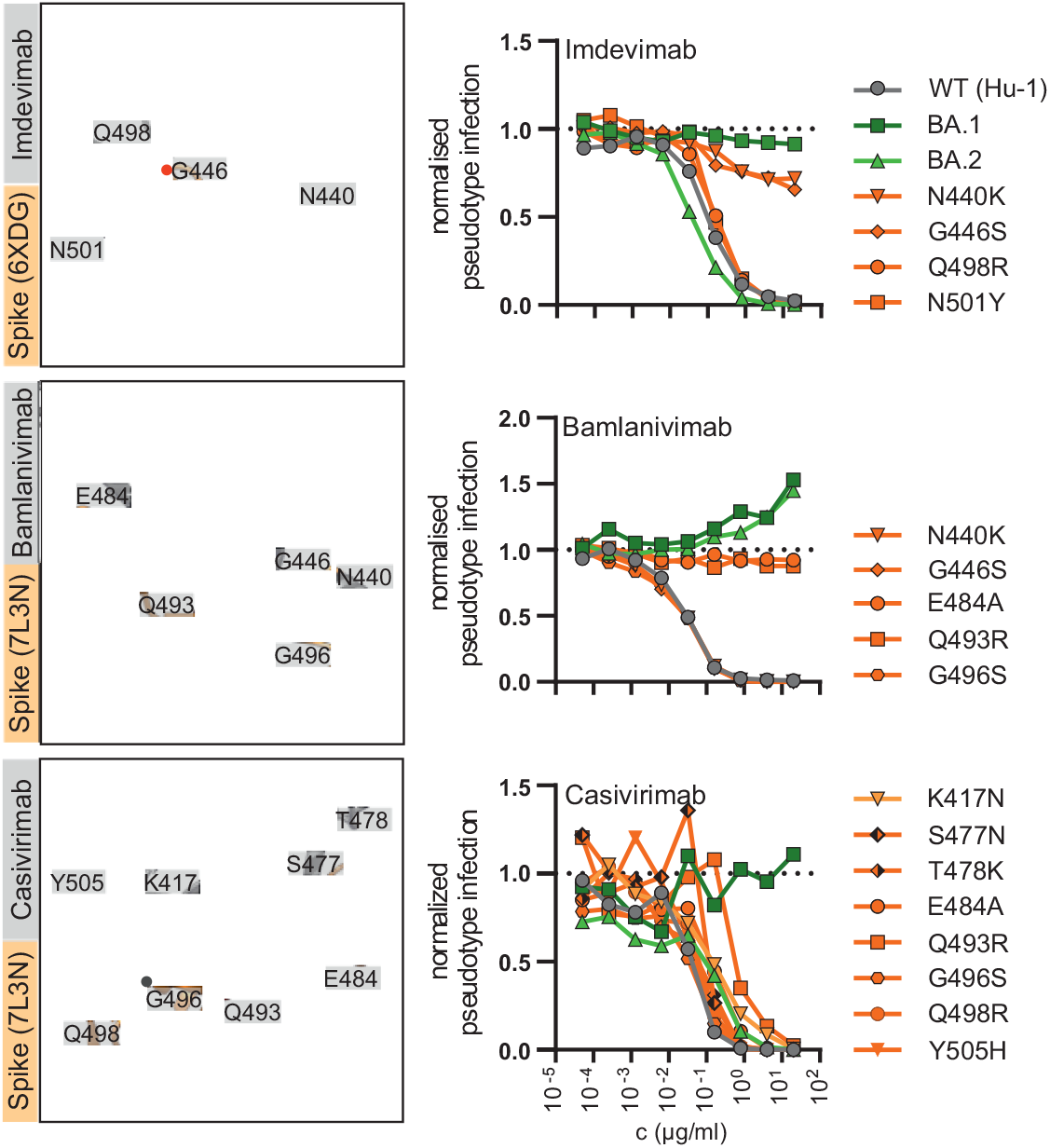
Impact of mutations in the Omicron Spike on neutralization by therapeutic Abs. Close-up view of neutralising antibodies binding the SARS-CoV-2 Spike (PDB: 6XDG or 7L3N as indicated) and automated quantification of GFP fluorescence of Caco-2 cells infected with VSVΔG-GFP pseudotyped with the indicated S variants. VSVpp were pre-treated (30 min, 37 °C) with the indicated amounts of Imdevimab, Bamlanivimab or Casivirimab. Lines represent the mean of three independent experiments.

## DISCUSSION

The Omicron VOC has outcompeted the previously dominating Delta VOC in an amazingly short time. It is generally accepted that the high number of changes in Omicron S are the main reason for effective spread of this VOC and confer increased transmission efficiency and escape from neutralizing antibodies. Here, we systematically analyzed the functional impact of all individual amino acid changes, deletions and insertions that are characteristic for the Omicron BA.1 and BA.2 VOCs. In total, we examined 48 mutant Spike constructs containing amino acid changes distinguishing BA.1 and BA.2 Omicron VOCs from the original Hu-1 Wuhan strain. We identified several changes that strongly impair Spike-mediated infection and proteolytic processing. In addition, we demonstrate that BA.1 or BA.2 specific mutations in the NTD as well as shared alterations in the RBD significantly reduce the susceptibility of Spike containing VSVpp to neutralization by sera from BNT/BNT vaccinated individuals and therapeutic antibodies.

One striking finding was that individual mutations of S371F/L, S375F, T376A and (to a lesser extent) S373P in the receptor binding domain strongly impair Spike-containing pseudoparticle infectivity and Spike processing. S375F had the most drastic effect and almost fully disrupted Spike function and processing. This agrees with a recent preprint (Yamasoba et al., 2022) and is of particular interest because it has recently been reported that the S371L and S371F mutations in BA.1 and BA2, respectively, have major effects on neutralization by different RBD classes (Liu et al., 2021; Iketani et al., 2022). However, our results show that although these mutations significantly reduce S-mediated infection (Figure 2) they have only modest effects on neutralization by BNT/BNT sera (Figure 6). Notably, all these residues are part of a loop that may affect the open and closed conformation of the RBD. Recent structural analyses show that mutations of S371L, S373P and S375F promote interprotomer interactions between the “down” RBDs (Gobeil et al., 2022). Specifically, it has been proposed that the S373P substitution induces conformational changes of the loop resulting in closer packing of the RBD-RBD interface via interactions of S373P and S375F with the N501Y and Y505H substitutions in the adjacent RBD (Gobeil et al., 2022). Our functional analyses show that mutations of S371L/F and T376A severely affect Spike function, while changes of N501Y and Y505H had no disruptive effects on S-mediated infection. Both individual and combined mutations in the three serine residues (S371, S373 and S375) severely impaired the ability of the Hu-1 Spike protein to mediate virus-cell and cell-cell fusion. While further studies are necessary, it is tempting to speculate that, in the absence of additional alterations, changes of S371L, S373P, S375F and perhaps T376A might stabilize the inactive closed conformation of the Spike protein. The severe disruptive effects also raise questions whether they are associated with a selection advantage. Our identification of a subcluster of BA.1 isolates that sequentially reverts to the original sequence (FxPxF to FxSxF to FxSxS to SxSxS; Figure 4A) indicates that this may not be the case and it will be of significant interest to closely monitor the frequency of Omicron variants containing alterations in the serine containing loop region.

A variety of mutations in the S1 subunit (ΔH69/V70, T95I, ΔY144, K417N, T478K, N501Y, D614G, H655Y and P681H) of BA.1 and/or BA.2 Omicron S proteins have previously been observed in other VOCs (Golcuk et al., 2021). As previously suggested (Mannar et al., 2022), we found that deletion of ΔH69/V70 increased S-mediated infectivity (Figure 2). Our analyses also confirmed (Lam et al., 2021) that mutation of N501Y in the RBM of the Omicron Spike increases the binding affinity to ACE2 (Figure 5A). It has been proposed that mutations of H655Y, N679K and P681H may increase furin-mediated S1/S2 cleavage and enhance pseudoparticle infectivity (Aggarwal et al., 2021; Cameroni et al., 2021; VanBlargan et al., 2022). We found that none of these three changes significantly enhanced S-mediated VSVpp infection (Figure 2) and only H655Y clearly enhanced S processing (Figure 3). Mutations of K417N, Q493R, Q498R and N501Y are identical or similar to changes emerging during SARS-CoV-2 adaptation to experimentally infected mice (Huang et al., 2021; Sun et al., 2021), and were proposed to stabilize the RBD and ACE2 interaction (Meng et al., 2022) or to contribute to the ability of Omicron to infect mouse cells (Hoffmann et al., 2021). However, individual changes had no significant effect on VSVpp infectivity or processing but reduced susceptibility to BNT/BNT neutralization. Similarly, based on cryo-EM analyses it has been suggested that alterations of Q493R, G496S and Q498R in the RBD of the Omicron S form may allow the formation of stronger interactions with ACE2 that compensate for a disruptive effect of K417N (Mannar et al., 2022) but none of these four mutations individually significantly affected S-mediated infection (Figure 2).

Predictably, many shared mutations in the RBD domain of BA.1 and BA.2 S proteins reduced the sensitivity of VSVpp to neutralization by sera from BNT/BNT vaccinated individuals (Figure 6A). In addition, we also found that mutations of N440K or G446S conferred resistance to imdevimab and changes of E484A or Q493R to bamlanivimab, respectively. This was expected since these mutations are located within the epitopes bound by these antibodies (Figure 6B, 6C). Our results add to the evidence (Iketani et al., 2022; Liu et al., 2022) that single amino acid changes may confer full resistance to neutralizing antibodies. In comparison, mutation of E484A that has also been observed in other SARS-CoV-2 VOCs and was suggested to be associated with immune-escape (Rath et al., 2022) had only marginal effects on neutralization sensitivity. Unexpectedly, most lineage-specific changes in the NTD, such as A27S, T95I, Δ142-144, G142D, INS214EPE, L212I and V213G, were at least as effective in reducing S sensitivity to neutralization by sera from BNT/BNT vaccinated individuals as changes in the RBD (Figure 6A). This supports a key role of the NTD as target for neutralizing antibodies in sera from vaccinated individuals.

Three mutations (Q954H, N969K and L981F) are located in the HR1 region of the S2 subunit of the S protein (Figure 1D). It has been initially proposed that these changes may promote 6-helix bundle formation and subsequent fusion (Sarkar et al., 2021) but more recent evidence suggests that they may attenuate rather than enhance S-mediated fusion efficiency (Suzuki et al., 2022). In agreement with the latter, changes of Q954H and N969K clearly reduced S-mediated VSVpp infection (Figure 2). In contrast, substitution of L981F enhanced Spike-mediated VSVpp infection and (more strongly) cell-to-cell fusion (Figure 5B). Reactive force simulations suggest that the L981F mutation enhances interactions between the HR1 and HR2 regions that drive fusion. Notably, recent data showed that the three mutations in the HR1 region of the Omicron S do not alter the global architecture of the post-fusion six-helix bundle (Yang et al., 2022) and peptide-based pan-CoV fusion inhibitors derived from the HR region maintain high potency against the SARS-CoV-2 Omicron VOC (Xia et al., 2022).

The molecular mechanisms of several mutations in the Omicron S protein remain to be fully elucidated. For example, BA.2-specific changes of T19I and Δ24-26 in the NTD severely reduced S-mediated infection and processing although they do not affect known functional domains. It has been suggested that a shared mutation of N764K and a BA.2-specific substitution of N856K generate potential cleavage sites for SKI-1/S1P protease and might impede the exposition of the fusion peptide for membrane fusion (Maaroufi, 2022). We found that N764K is indeed associated with increased infectivity and increased levels of processed Spike in VSVpp. In comparison, N856K clearly reduced S-mediated infection despite normal processing.

### Limitations of the study

In the present study, we used pseudotyped viral particles instead of replication-competent recombinant SARS-CoV-2 variants, which serves as a proxy to assess infectivity, fusion activity and incorporation. In addition, the impact of many changes might be context-dependent and this might explain why some individual changes had disruptive effects on Hu-1 S function although they are found in Omicron S proteins. It is difficult to predict which of the numerous mutations in the Omicron S might compensate for disruptive mutations. For example, we found that the BA.1 S was less effective in mediating infection than the BA.2 S protein (Figure 2). This agrees, with accumulating evidence that the BA.2 VOC might be more infectious and more virulent than the BA.1 VOC (Suzuki et al., 2022). In addition, we analysed only a limited number of sera from individuals who received a single vaccine regimen (BNT/BNT) and just a few therapeutic antibodies. While further studies are required to fully understand the full consequences of all the complex changes in the Omicron Spike on viral infectivity, tropism, transmission and pathogenesis our results provide first important insights into the functional impact of mutations characteristic for the Omicron VOC Spike that currently dominates the pandemic.

## Supporting information

Suppemental File 1

## ACKNOWLEDGEMENTS

We thank Kerstin Regensburger, Regina Burger, Jana-Romana Fischer, Birgit Ott, Martha Meyer, Nicola Schrott and Daniela Krnavek for technical assistance and Dorota Kmiec for critical reading of our manuscript. The ACE2 expression vector and SARS-CoV-2 S-HA plasmid were kindly provided by Shinji Makino and Stefan Pöhlmann. C.P. and S.N. are part of the International Graduate school for Molecular Medicine (IGradU). This study was supported by DFG grants to F.K. (CRC 1279, SPP 1923), K.M.J.S. (CRC 1279, SPP 1923, SP 1600/6-1) and T.J. (CRC 1279). F.K., and K.M.J.S. were supported by the BMBF (Restrict SARS-CoV-2 and IMMUNOMOD).

## CONTRIBUTIONS

C.P. and F.Z. performed most experiments with support by S.N. F.Z., K.M.J.S., and F.K. conceived the study and planned experiments. C.J. and T.J. performed molecular modelling and dynamics simulations. F.K. wrote the initial draft of the manuscript. All authors reviewed and approved the manuscript.

## STAR METHODS

### RESOURCE AVAILABILITY

#### Lead contact

Further information and requests for resources and reagents should be directed to and will be fulfilled by the Lead Contact, Frank Kirchhoff

#### Materials Availability

All unique reagents generated in this study are listed in the Key Resources Table and available from the Lead Contact.

#### Data and code availability

This study did not generate or analyse datasets or codes.

### METHOD DETAILS

#### Cell Culture

All cells were cultured at 37 °C and 5% CO_2_ in a humified atmosphere. HEK293T (human embryonic kidney) cells (ATCC: #CRL3216) were maintained in Dulbecco’s Modified Eagle Medium supplemented with 10% (v/v) heat-inactivated fetal calf serum, 2 mM L-glutamine, 100 μg/ml streptomycin and 100 U/ml penicillin. Caco-2 (human epithelial colorectal adenocarcinoma) cells were cultivated in DMEM containing 10% FCS, 2mM glutamine, 100 μg/ml streptomycin and 100 U/ml penicillin, 1mM NEAA supplement. Mouse I1-Hybridoma cells (ATCC: #CRL2700) were cultured in Roswell Park Memorial Institute (RPMI) 1640 medium supplemented with 10% (v/v) heat-inactivated fetal calf serum, 2 mM L-glutamine, 100 μg/ml streptomycin and 100 U/ml penicillin.

#### Expression Constructs

pCG_SARS-CoV-2-Spike-IRES_eGFP encoding the spike protein of SARS-CoV-2 isolate Wuhan-Hu-1(NCBI reference Sequence YP_009724390.1), pCG_SARS-CoV-2-S (1.617), pCG1_SARS-2-S (B.1.1.529) and pCG1_SARS-2-SΔ18 (BA.2) were kindly provided by Stefan Pöhlmann (DPZ Göppingen, Germany). pCG_SARS-CoV-2-Spike C-V5-IRES_eGFP was PCR amplified and subcloned into a pCG-IRES_eGFP expression construct using the restriction enzymes XbaI+MluI. The SARS-CoV-2 S mutant plasmids were generated using Q5 Site-Directed Mutagenesis Kit (NEB #E0554). ACE2 was synthezised by Twist bioscience, PCR amplified, and subcloned into a pCG-IRES_eGFP expression construct using the restriction enzymes XbaI+MluI. All constructs were verified by sequence analysis using a commercial sequencing service (Eurofins Genomics).

#### Molecular dynamics simulation

Initial atomic positions of ACE2-bound to SARS-CoV-2 spike (7KNB, https://www.rcsb.org/structure/7KNB) respectably the post-fusion structure of SARS-CoV-2 spike glycoprotein (PDB id 6M3W https://www.rcsb.org/structure/6m3w) were obtained from the Protein Data Bank (Bernstein et al., 1977). Equilibrations (300 K for 0.5 ns) were performed by ReaxFF (reactive molecular dynamic) simulations (Adri C. T. van Duin et al., 2001) within the Amsterdam Modelling Suite 2020 (http://www.scm.com). Based on the equilibrated structures, amino acids from the Wuhan-1 spike protein were replaced with the respective amino acids from Omicron BA.1 and BA.2 spike protein. These modified structures were additionally equilibrated (300 K for 0.5 ns) ReaxFF (reactive molecular dynamic) within an NVT while coupling the system to a Berendsen heat bath (T = 300 K with a coupling constant of 100 fs). The interaction energies were obtained by averaging over the last 0.1 ns of these simulations. The Visual Molecular Dynamics program (VMD 1.9.3) (Humphrey et al., 1996) was used for all visualizations.

#### Pseudoparticle Production

To produce pseudotyped VSV(GFP)ΔG particles, HEK293T cells were transfected with Spike-expressing vectors using polyethyleneimine (PEI 1 mg/ml in H_2_O). Twenty-four hours post-transfection, the cells were infected with VSVΔG(GFP)*VSV-G at a MOI of 3. The inoculum was removed 1hour post-infection. Pseudotyped VSVΔG-GFP particles were harvested 16 h post-infection. Remaining cell debris were removed by centrifugation (500 × g for 5 min). Residual particles carrying VSV-G were blocked by adding 10% (v/v) of I1-Hybridoma supernatant (I1, mouse hybridoma supernatant from CRL-2700; ATCC) to the cell culture supernatant.

#### Infection Assay

Caco-2 cells were infected with 100 μl of VSVΔG-GFP pseudo particles in 96 well format. GFP-positive cells were automatically counted using a Cytation 3 microplate reader (BioTek Instruments).

#### Pseudoparticle inhibition

50 μl of VSVΔG-GFP pseudo particles were preincubated for 30 min at RT with the indicated amounts of monoclonal antibodies (Bamlanivimab, Imdevimab, Casivirimab) or sera from fully BNT162b2 vaccinated individuals and transduced on CaCo-2 cells in 96 well format. 24 hours after infection, GFP-positive cells were automatically counted by a Cytation 3 microplate reader (BioTek Instruments).

#### Sera from vaccinated individuals

Blood samples of fully BNT162b2 vaccinated individuals were obtained after the participants information and written consent. Samples were collected 13–30 days after the second vaccination using S-Monovette Serum Gel tubes (Sarstedt). Before use, the serum was heat-treated at 56 °C for 30 min. Ethics approval was provided by the Ethic Committee of Ulm University (vote 99/21–FSt/Sta).

#### Whole-cell and cell-free lysates

To prepare whole-cell lysates, cells were collected and washed in phosphate-buffered Saline (PBS), pelleted and lysed in transmembrane lysis buffer, containing protease inhibitor (1:500). After 5 min of incubation on ice, supernatants were cleared by centrifugation (4 °C, 20 min, 20,817 × g). To prepare WB lysates of viral particles, the supernatants were layered on a cushion of 20% sucrose and centrifuged (4 °C, 90 min, 20,817 × g). The virus pellet was lysed in transmembrane lysis buffer, mixed with 4x Protein Sample Loading Buffer (LI-COR) containing 10% β-mercaptoethanol (Sigma Aldrich) and denaturized at 95 °C for 10 min.

#### SDS-PAGE and immunoblotting

SDS-PAGE and immunoblotting was performed as previously described (Zech et al. 2021). In brief, whole cell lysates were mixed with 4x Protein Sample Loading Buffer (LI-COR) containing 10% β-mercaptoethanol (Sigma Aldrich), heated at 95 °C for 20 min, separated on NuPAGE 4-12% Bis-Tris Gels (Invitrogen) for 90 min at 120 V and blotted at constant 30 V for 30 min onto Immobilon-FL PVDF membrane. After the transfer, the membrane was blocked in 1% Casein in PBS. Proteins were stained using primary antibodies directed against rabbit anti-V5 (Cell Signaling #13202; 1:1000), VSV-M (Absolute Antibody, 23H12, #Ab01404-2.0; 1:2000), actin (Anti-beta Actin antibody Abcam, ab8227, 1:5000,) and Infrared Dye labeled secondary antibodies (LI-COR IRDye) IRDye 800CW Goat anti-Mouse #926-32210, IRDye 680CW Goat anti-Rabbit (#925-68071), all 1:20,000. Proteins were detected using a LI-COR Odyssey scanner and band intensities were quantified using LI-COR Image Studio version 5.

#### ACE2 interaction assay

HEK293T cells expressing Spike were collected 48 h after the transfection, washed with phosphate-buffered saline (PBS), lysed in a non-denaturizing lysis buffer. Interaction between Spike protein and ACE2 was assessed through a Spike-ACE2 binding assay kit (COVID-19 Spike-ACE2 binding assay II, Ray Bio). Briefly, 10 μl of WCLs were diluted 1:5 in 1x assay diluent buffer (RayBio), added to ACE2 coated wells (RayBio) and incubated for 2 h with shaking. After washing extensively with the provided wash buffer (RayBio, #EL-ITEMB), the wells were incubated 1 h with 100 μl anti-V5(MS) (1:1,000, Cell Signalling, #80076), washed and incubated for 1 h with 100 μl anti-MS-HRP (1:1,000, RayBio). After washing, the samples were incubated with 50 μl of TMB Substrate Solution (RayBio, #EL-TMB) for 30 min. The reaction was stopped by the addition of 50 μl Stop Solution (RayBio, #ELSTOP) and absorbance was measured at 450 nm with a baseline correction at 650 nm.

#### Immunofluorescence

HEK293T cells were plated in 12-well tissue culture dishes on 13-mm round borosilicate cover slips pre-coated with poly-L-lysine. 24 hours after, the cells were transfected with expression constructs for Spike protein using polyethyleneimine (PEI 1 mg/ml in H_2_O). 24 hours after transfection, cells were washed with cold PBS and fixed in 4% paraformaldehyde solution (PFA) for 20 min at RT, permeabilized and blocked in PBS containing 0.5% Triton X-100 and 5% FCS for 30 min at RT. Thereafter, samples were washed with PBS and incubated for 2 h at 4°C with primary antibody (anti-V5(MS) (1:1,000, Cell Signalling, #80076)) diluted in PBS. The samples were washed with PBS/0.1% Tween 20 and incubated in the dark for 2 h at 4°C with the secondary antibody (Alexa Fluor-647-conjugated anti-mouse antibody, 1:1000, Thermo Fisher Scientific) and 500 ng/ml DAPI. After washing with PBS-T and water, cover slips were mounted on microscopy slides. Images were acquired using a Zeiss LSM800 confocal laser scanning microscope with ZEN imaging software (Zeiss).

#### Quantification of syncytia formation

To detect formation of syncytia, HEK293T cells were co-transfected with ACE2 and Spike expressing vectors using polyethyleneimine (PEI 1 mg/ml in H_2_O). Twenty-four hours post-transfection, fluorescence microscopy images were acquired using the Cytation 3 microplate reader (BioTek Instruments) and the GFP area was quantified using ImageJ.

#### Statistical analysis

Statistical analyses were performed using GraphPad PRISM 9.2 (GraphPad Software). P-values were determined using two-tailed Student’s t-test with Welch’s correction or One-Way ANOVA with multiple comparison against the Wuhan-Hu-1 values. Unless otherwise stated, data are shown as the mean of at least three independent experiments ± SEM. Significant differences are indicated as *p <0.05; **p <0.01; ***p <0.001.

##### Key resources table

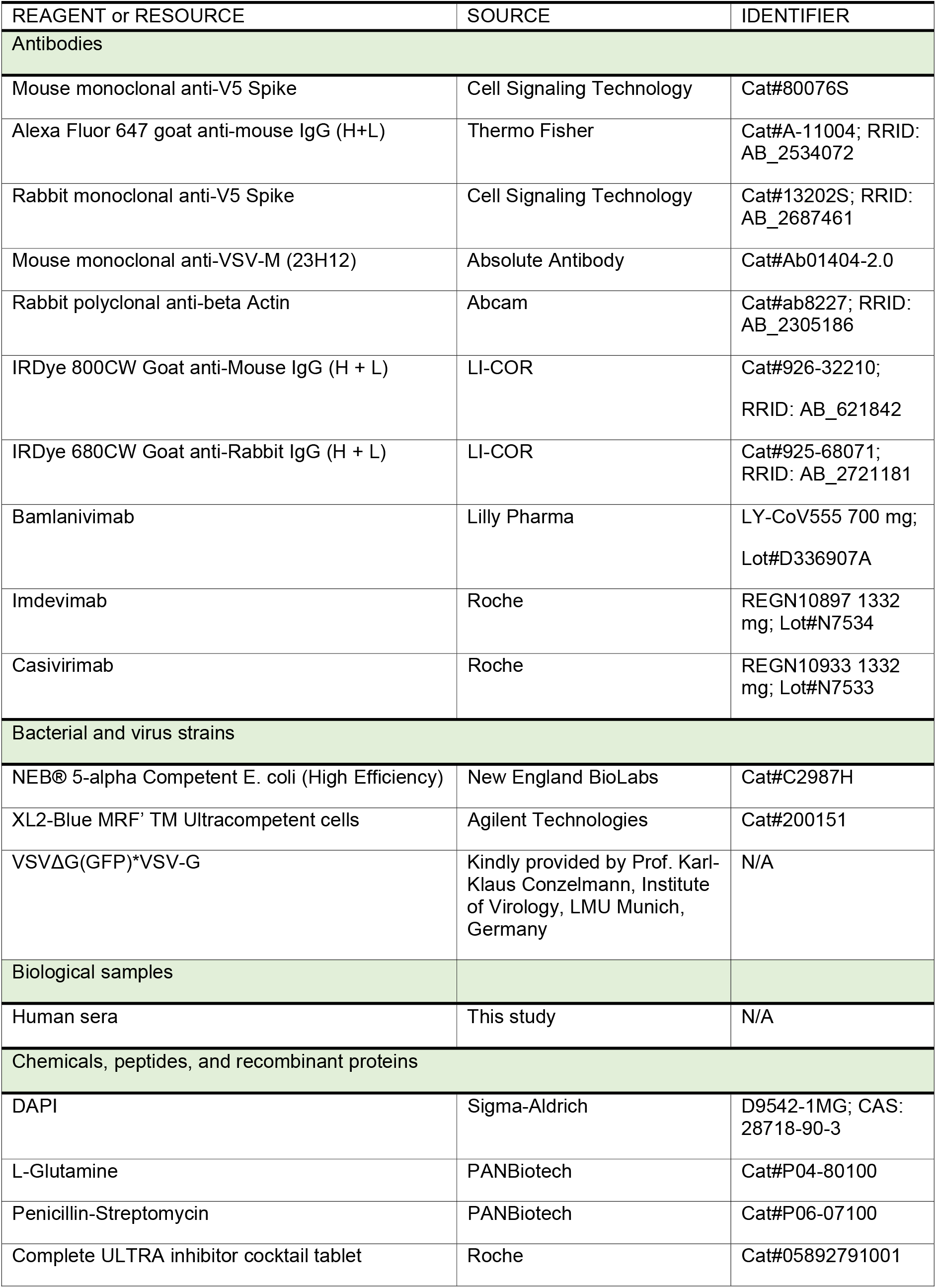
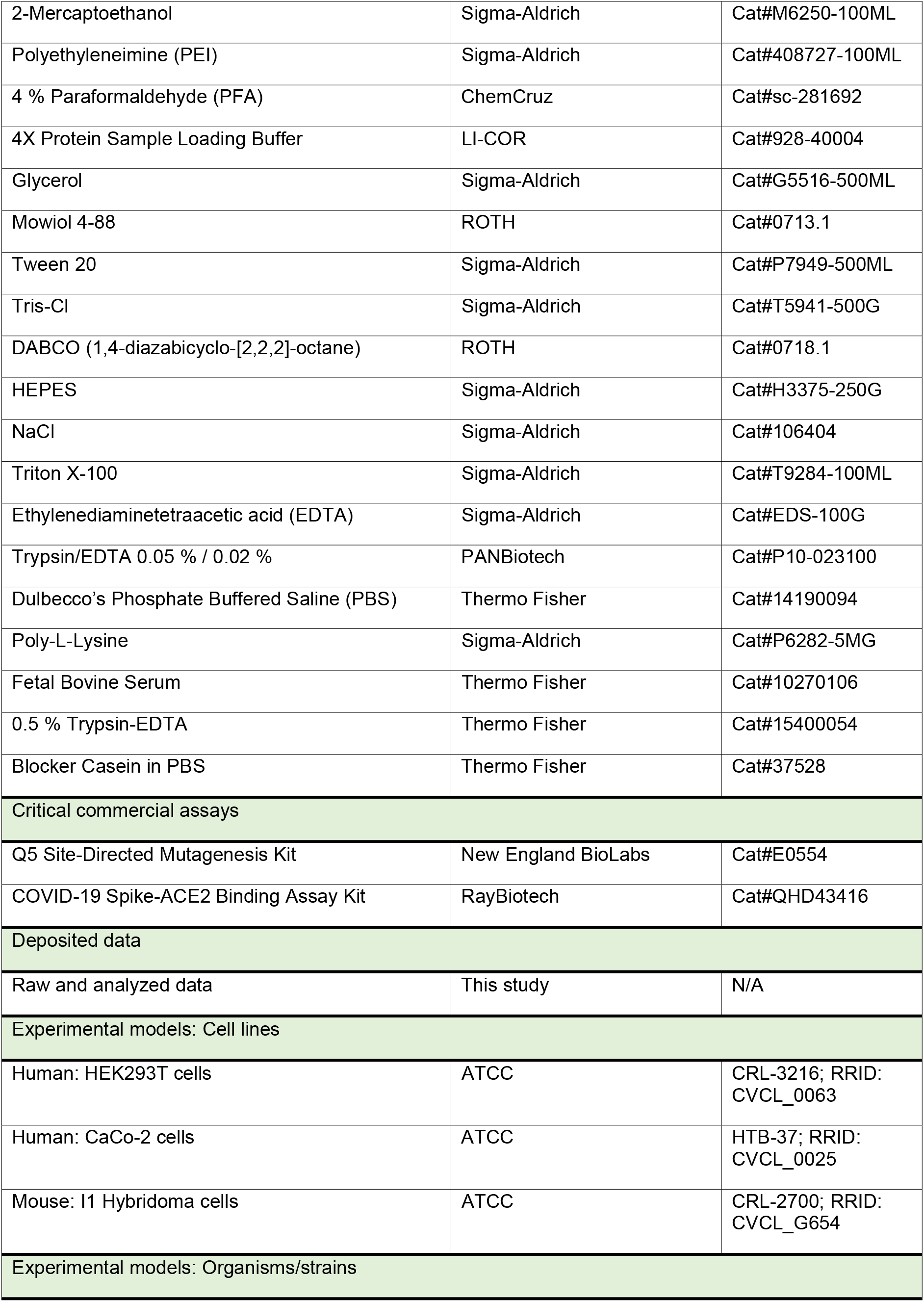
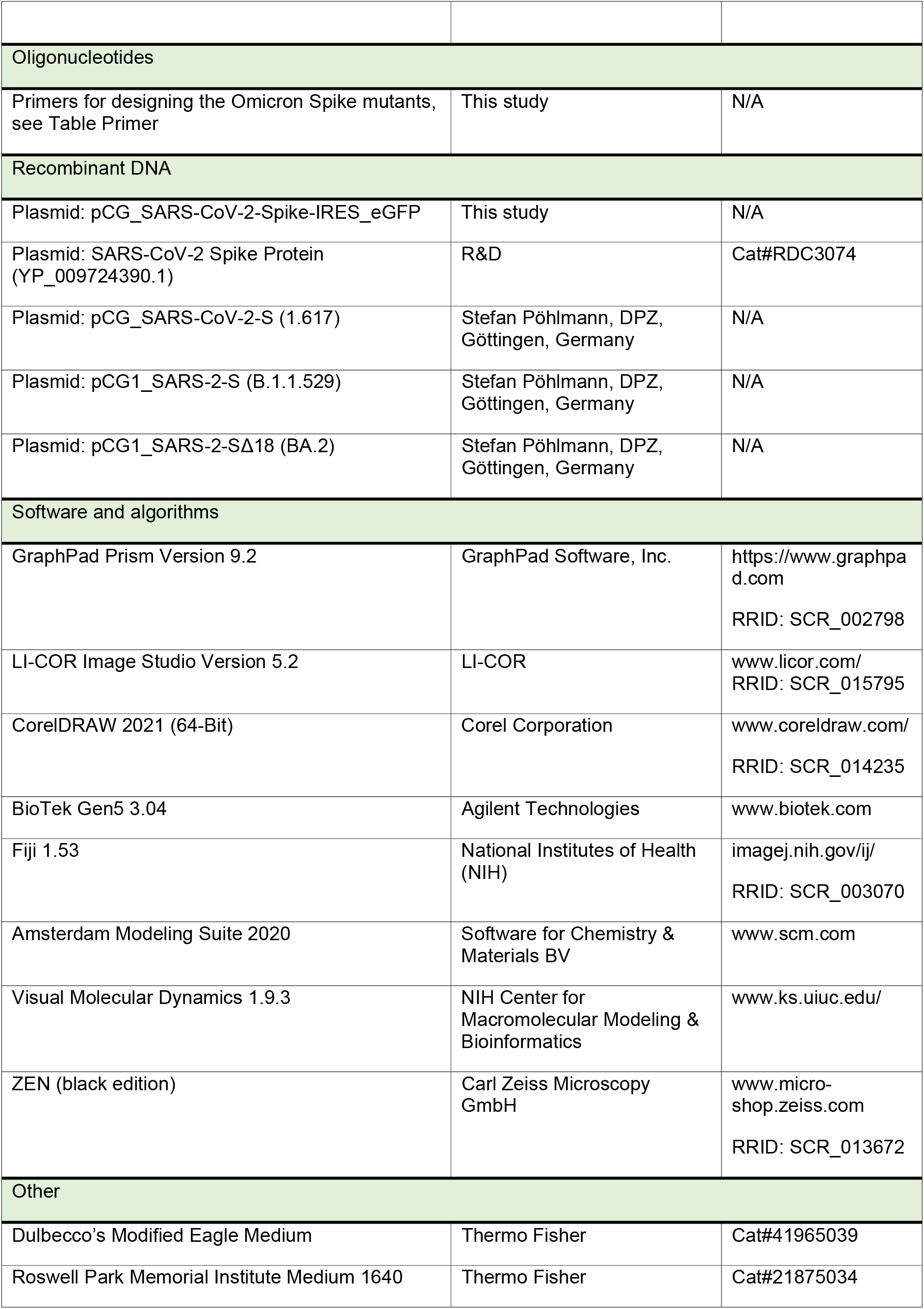
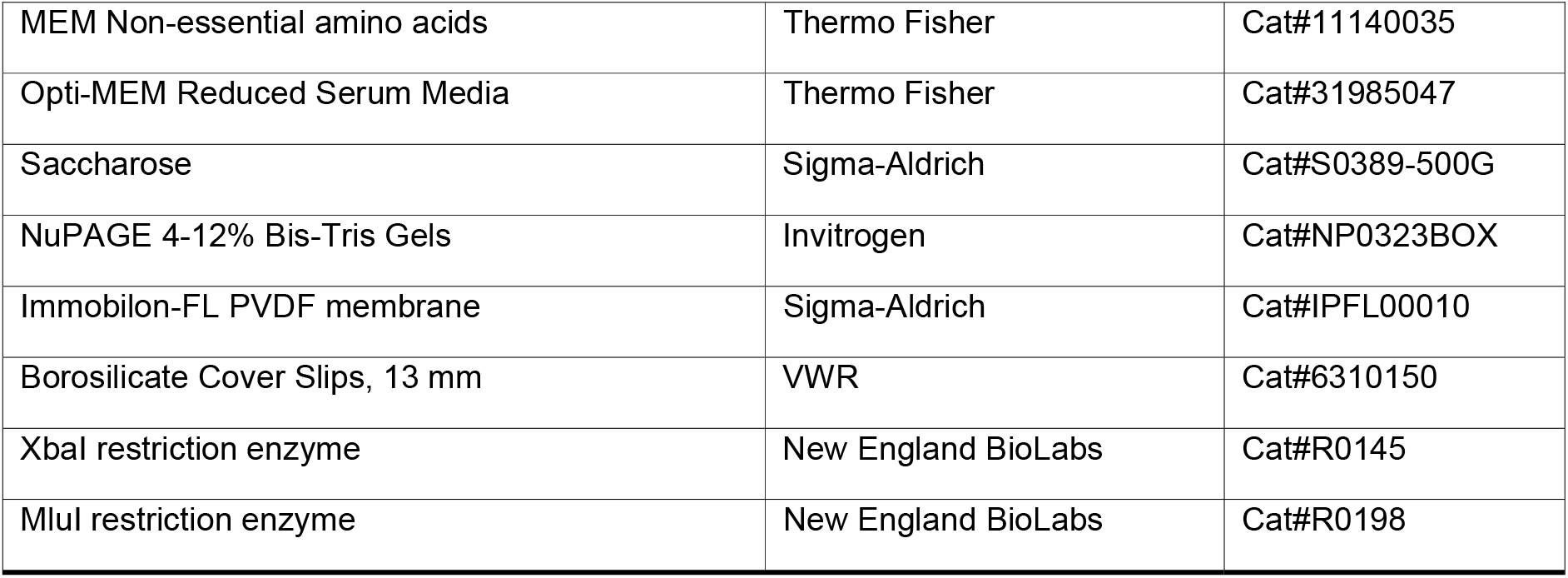

