## Supplementary material for "Determinants of Spike Infectivity, Processing and Neutralization in SARS-CoV-2 Omicron subvariants BA.1 and BA.2": Suppemental File 1

### SUPPLEMENTAL FIGURES

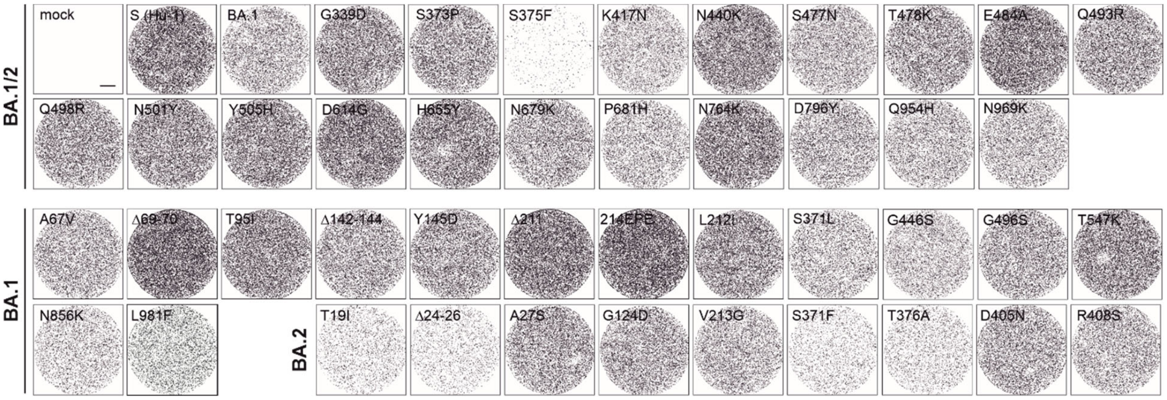


**Figure S1 (related to Figure 2). Infection of CaCo-2 cells by VSVpp containing WT or mutant S proteins.**

Images of CaCo-2 cells transduced with VSVΔG-GFP pseudotyped with the Hu-1 or mutant SARS-CoV-2 S proteins. Successful infection events (=GFP positive cells) are displayed as black dots. Scale bar, 1.5 μm.

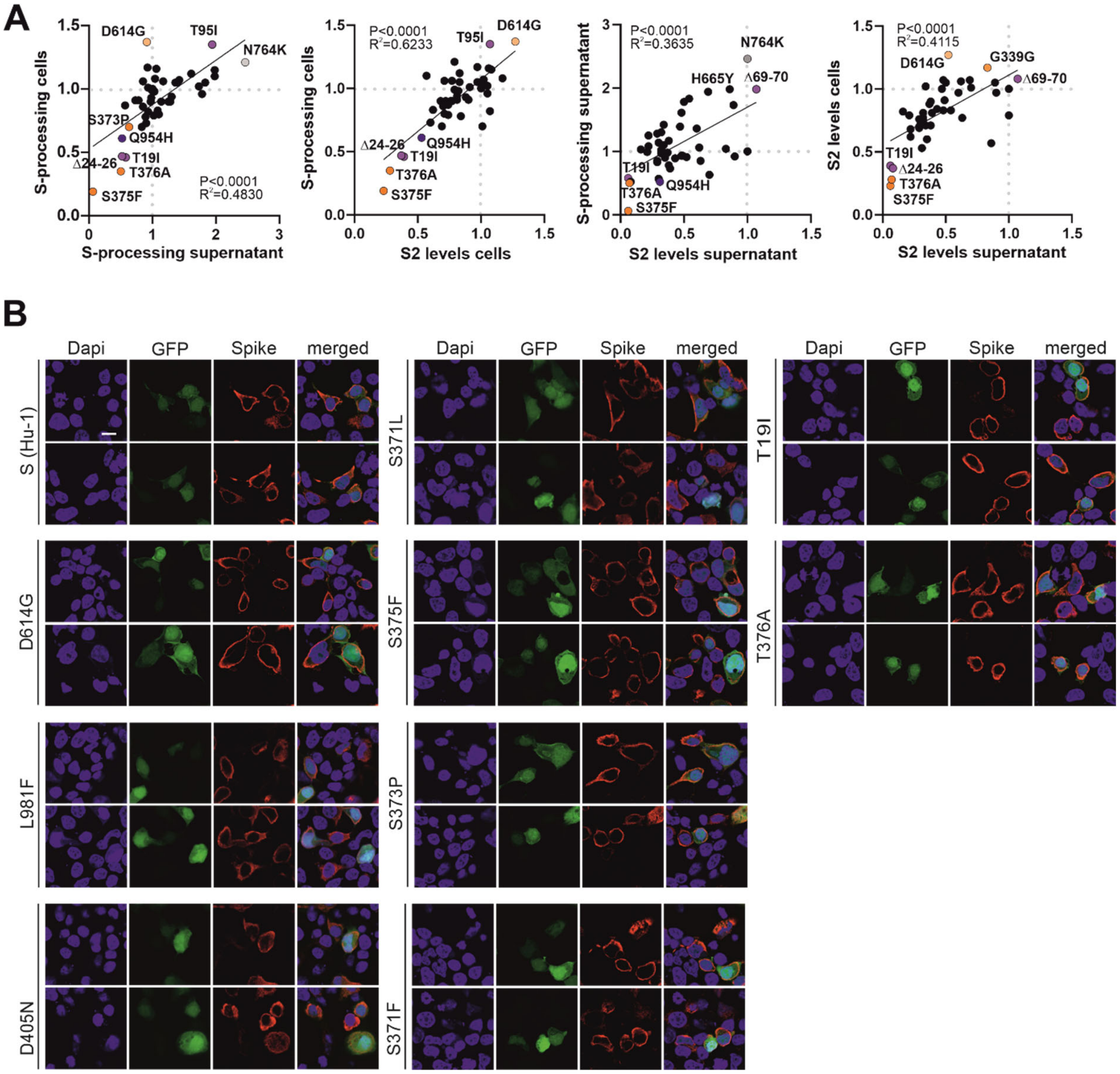


**Figure S2 (related to Figure 3). Correlation analyses and localization of S proteins.**

(A) Correlation of the between the indicated parameters. S2 expression levels and S/S2 processing of mutant S proteins were normalized to the parental Hu-1 S (set to 1).

(B) Immunofluorescence images of HEK293T cells expressing the parental Hu-1 or indicated mutant S proteins. Scale bar, 10 µm.

**Table S1. Origin and neutralizing activity of sera from BNT/BNT vaccinated individuals.**

| Donor | AGE | Sex |
|---|---|---|
| 1 | 61 | w |
| 2 | 28 | w |
| 3 | 58 | m |
| 4 | 27 | w |
| 5 | 37 | m |

**WT**

| Donor | S Hu-1 | Delta | BA.1 | BA.2 |
|---|---|---|---|---|
| 1 | 0,15 | 0,69 | 3,24 | 0,46 |
| 2 | 0,9 | 1,52 | >10 | 1,39 |
| 3 | 0,4 | 7,11 | >10 | >10 |
| 4 | 0,99 | 3,36 | >10 | 8,03 |
| 5 | 0,29 | 0,82 | 4,44 | 0,91 |

**Colors:**

| |
|---|
| NTD |
| RBD |
| RBM |
| S1 |
| S2 |
| S2' |

**IC50 Values BA.1/BA.2**

| Donor | G339D | S373P | S375F | K417N | N440K | S477N | T478K | E484A | Q493R | Q498R | N501Y | Y505H | D614G | H655Y | N679K | P681H | N764K | D796Y | Q954H | N969K |
|---|---|---|---|---|---|---|---|---|---|---|---|---|---|---|---|---|---|---|---|---|
| 1 | 0,29 | 0,88 | 0,17 | 1,23 | 1,09 | 0,61 | 0,46 | 0,51 | 0,89 | 0,6 | 0,84 | 0,85 | 0,69 | 0,29 | 0,77 | 0,68 | 0,77 | 0,28 | 0,17 | 0,32 |
| 2 | 1,07 | 0,97 | 0,29 | 0,87 | 0,73 | 0,55 | 0,12 | 0,2 | 0,34 | 0,55 | 0,41 | 0,43 | 0,58 | 0,43 | 0,41 | 0,62 | 0,41 | 0,19 | 0,15 | 0,33 |
| 3 | 1,1 | 1,82 | 0,31 | 2,79 | 2,15 | 1,28 | 2,9 | 0,78 | 1,86 | 1,54 | 2,53 | 1,77 | 1,45 | 0,82 | 1,17 | 0,9 | 1,42 | 0,45 | 0,42 | 0,57 |
| 4 | 6,73 | 9,11 | 4,43 | 4,4 | 5,33 | 2,49 | 0,1 | 1,14 | 6,07 | 2,57 | 3,63 | 3,35 | 3,23 | 2,16 | 3,8 | 2,88 | 2,02 | 0,63 | 0,95 | 1,45 |
| 5 | 0,34 | 0,42 | 0,16 | 0,4 | 0,27 | 0,22 | 0,1 | 0,07 | 0,31 | 0,22 | 0,27 | 0,21 | 0,22 | 0,2 | 0,24 | 0,42 | 0,31 | 0,07 | 0,13 | 0,21 |

**IC50 Values BA.1**

| Donor | A67V | d69-70 | T95I | d142-144 | Y145D | d211 | 214EPE | L212I | S371L | G446S | G496S | T547K | N856K | L981F |
|---|---|---|---|---|---|---|---|---|---|---|---|---|---|---|
| 1 | 0,01 | 0,12 | 0,31 | 0,49 | 1,32 | 0,01 | 1,47 | 1,03 | 0,84 | 0,16 | 0,81 | 0,33 | 0,32 | 0,38 |
| 2 | 0,26 | 0,27 | 0,79 | 1,09 | 1,01 | 0,06 | 2,22 | 1,04 | 0,64 | 0,28 | 0,74 | 0,39 | 0,21 | 0,24 |
| 3 | 0,04 | 0,19 | 2,59 | 4,61 | 5,18 | 0,02 | 2,92 | 3,48 | 1,84 | 1,03 | 1,7 | 0,85 | 0,59 | 0,37 |
| 4 | 0,55 | 0,84 | 1,41 | 2,64 | 9,56 | 0,03 | 9,89 | 7,56 | 3,91 | 1,55 | 3,77 | 1,49 | 1,83 | 0,72 |
| 5 | 0,04 | 0,05 | 0,5 | 0,68 | 0,38 | 0,05 | 0,64 | 0,6 | 0,15 | 0,21 | 0,29 | 0,23 | 0,07 | 0,1 |

**IC50 Values BA.2**

| Donor | T19I | d24-26 | A27S | G142D | V213G | S371F | T376A | D405N | R408S |
|---|---|---|---|---|---|---|---|---|---|
| 1 | 0,16 | 0 | 1,94 | 0,43 | 0,31 | 0,36 | 0,39 | 0,04 | 1,25 |
| 2 | 0,22 | 1,08 | 0,96 | 0,92 | 0,93 | 0,21 | 0,5 | 0,44 | 1,96 |
| 3 | 0,75 | 0,11 | 1,92 | 5,88 | 2,74 | 2,66 | 1,17 | 0,06 | 2,96 |
| 4 | 2,51 | 8,79 | 7,39 | 2,18 | 9,74 | 9,24 | 3,88 | 1,64 | 6,73 |
| 5 | 0,15 | 0,25 | 0,56 | 0,53 | 0,55 | 0,27 | 0,12 | 1,53 | 0,52 |

**Table S2. Primers used for site-directed mutagenesis of the Hu-1 Spike.**

| Vector | Mutation at amino acid position | Primer Sequence |
|---|---|---|
| pCG_SARS-CoV-2-Spike C-V5-IRES_eGFP | **T19I_F** | GTGAACCTGAtCACAAGAACCC |
| | **T19I_R** | ACACTGGCTGGACACCAG |
| | **Δ24-26-F** | GCCTACACCAACAGCTTT |
| | **Δ24-26-R** | CTGGGTTCTTGTGGTCAG |
| | **A27S_F** | GCTGCCTCCAtCCTACACCAA |
| | **A27S_R** | TGGGTTCTTGTGGTCAGG |
| | **A67V_F** | TGGTTCCACGtCATCCACGTG |
| | **A67V_R** | GGTCACGTTGCTGAAGAAAG |
| | **Δ69-70-F** | TCCGGCACCAATGGCACC |
| | **Δ69-70-R** | GATGGCGTGGAACCAGGTC |
| | **T95I_F** | TTTGCCAGCAtCGAGAAGTCC |
| | **T95I_R** | GTACACCCCGTCGTTGAAG |
| | **Δ142-144-F** | TATCACAAGAACAACAAGAGCTGG |
| | **Δ142-144-R** | CAGGAAGGGGTCGTTGCA |
| | **G142D_F** | CCCTTCCTGGaCGTCTACTATC |
| | **G142D_R** | GTCGTTGCAGAACTGGAAC |
| | **Y145D_F** | GGGCGTCTACgATCACAAGAAC |
| | **Y145D_R** | AGGAAGGGGTCGTTGCAG |
| | **Δ211-F** | CTCGTGCGGGATCTGCCT |
| | **Δ211-R** | GATAGGGGTGTGCTTGCTG |
| | **214EPE_F** | cgagGATCTGCCTCAGGGCTTC |
| | **214EPE_R** | ggctcCCGCACGAGGTTGATAGG |
| | **L212I_F** | CCCTATCAACaTCGTGCGGGA |
| | **L212I_R** | GTGTGCTTGCTGTAGATCTTG |
| | **V213G_F** | ATCAACCTCGgGCGGGATCTG |
| | **V213G_R** | AGGGGTGTGCTTGCTGTAG |
| | **G339D_F** | TGCCCCTTCGaCGAGGTGTTC |
| | **G339D_R** | CAGATTGGTGATATTGGGGAAC |
| | **S371L_F** | GCTGTACAACctCGCCAGCTTCAGCAC |
| | **S371L_R** | ACGGAGTAGTCGGCCACG |
| | **S371F_F** | CTGTACAACTtCGCCAGCTTC |
| | **S371F_R** | CACGGAGTAGTCGGCCAC |
| | **S373P_F** | CAACTCCGCCccCTTCAGCACC |
| | **S373P_R** | TACAGCACGGAGTAGTCG |
| | **S375F_F** | CGCCAGCTTCttCACCTTCAAG |
| | **S375F_R** | GAGTTGTACAGCACGGAG |
| | **T376A_F** | CAGCTTCAGCgCCTTCAAGTG |
| | **T376A_R** | GCGGAGTTGTACAGCACG |
| | **D405N_F** | GATCCGGGGAaATGAAGTGCG |
| | **D405N_R** | ACGAAGCTGTCGGCGTAC |
| | **R408S_F** | AGATGAAGTGtcGCAGATTGCCCCTGG |
| | **R408S_R** | CCCCGGATCACGAAGCTG |
| | **K417N_F** | AGACAGGCAACcATCGCCGACTAC |
| | **K417N_R** | GTCCAGGGGCAATCTGCC |
| | **N440K_F** | ACAGCAACAAgCTGGACTCCA |
| | **N440K_R** | TCCAGGCAATCACACAGC |
| | **G446S_F** | CTCCAAAGTCaGCGGCAACTAC |
| | **G446S_R** | TCCAGGTTGTTGCTGTTC |
| | **S477N_F** | CAGGCCGGCAaCACCCCTTGT |
| | **S477N_R** | ATAGATCTCGGTGGAGATGTCCCG |
| | **T478K_F** | GCCGGCAGCAaaCCTTGTAACG |
| | **T478K_R** | CTGATAGATCTCGGTGGAG |
| | **E484A_F** | AACGGCGTGGcAGGCTTCAACTGC |
| | **E484A_R** | ACAAGGGGTGCTGCCGGC |
| | **Q493R_F** | TTCCCACTGCgGTCCTACGGC |
| | **Q493R_R** | GTAGCAGTTGAAGCCTTCCAC |
| | **G496S_F** | GCAGTCCTACaGCTTTCAGCC |
| | **G496S_R** | AGTGGGAAGTAGCAGTTG |
| | **Q498R_F** | TACGGCTTTCgGCCCACAAAT |
| | **Q498R_R** | GGACTGCAGTGGGAAGTAG |
| | **N501Y_F** | TCAGCCCACAtATGGCGTGGG |
| | **N501Y_R** | AAGCCGTAGGACTGCAGTG |
| | **Y505H_F** | TGGCGTGGGCcATCAGCCCTA |
| | **Y505H_R** | TTTGTGGGCTGAAAGCCGTAG |
| | **T547K_F** | AACGGCCTGAaaGGCACCGGCG |
| | **T547K_R** | GAAGTTGAAGTTCACGCATTTG |
| | **D614G_F** | CTGTACCAGGgCGTGAACTGT |
| | **D614G_R** | CACTGCCACCTGATTGCT |
| | **H655Y_F** | CGGAGCCGAGtACGTGAACAA |
| | **H655Y_R** | ATCAGACAGCCGGCTCTG |
| | **N679K_F** | CACAGACAAAgAGCCCCAGAC |
| | **N679K_R** | TCTGGTAGCTGGCACAGA |
| | **P681H_F** | ACAAACAGCCaCAGACGGGCC |
| | **P681H_R** | CTGTGTCTGGTAGCTGGCAC |
| | **N764K_F** | CCCAGCTGAAgAGAGCCCTGA |
| | **N764K_R** | TGCAGAAGCTGCCGTACT |
| | **D796Y_F** | TCCTATCAAGtACTTCGGCGG |
| | **D796Y_F** | GGGGTCTTGTAGATCTGC |
| | **N856K_F** | AGAAGTTTAAgGGACTGACAGTGC |
| | **N856K_R** | GGGCGCAAATCAGATCCC |
| | **Q954H_F** | TGGTCAACCAcAATGCCCAGG |
| | **Q954H_R** | CGTCCTGCAGCTTTCCCA |
| | **N969K_F** | TGTCCTCCAAgTTCGGCGCCA |
| | **N969K_R** | GCTGCTTGACCAGGGTGTTC |
| | **L981F_F** | GAACGATATCtttAGCAGACTGGACAAGG |
| | **L981F_R** | AGCACAGAGCTGATGGCG |
| | **S371L/S373P/S375F_F** | CAACCTCGCCcccCTTCTTCACCTTC |
| | **S371L/S373P/S375F_R** | TACAGCACGGAGTAGTCG |
| | **S371F/S373P/S375F_F** | CAACTTCGCCcccCTTCTTCACCTTC |
| | **S371F/S373P/S375F_R** | TACAGCACGGAGTAGTCG |
| | **S371F/S373P/S375F_R** | AAGTTGTACAGCACGGAG |
